# Stability and similarity of the pediatric connectome as developmental measures

**DOI:** 10.1101/828137

**Authors:** Tamara Vanderwal, Jeffrey Eilbott, Clare Kelly, Simon R. Frew, Todd S. Woodward, Michael P. Milham, F. Xavier Castellanos

## Abstract

Patterns of functional connectivity are unique at the individual level, enabling test-retest matching algorithms to identify a subject from among a group using only their functional connectome. Recent findings show that accuracies of these algorithms in children increase with age. Relatedly, the persistence of functional connectivity (FC) patterns across tasks and rest also increases with age. This study investigated the hypothesis that within-subject stability and between-subject similarity of the whole-brain pediatric connectome are developmentally relevant outcomes. Using data from 210 help-seeking children and adolescents, ages 6-21 years (Healthy Brain Network Biobank), we computed whole-brain FC matrices for each participant during two different movies (MovieDM and MovieTP) and two runs of task-free rest (all from a single scan session) and fed these matrices to a test-retest matching algorithm. We replicated the finding that matching accuracies for children and youth (ages 6-21 years) are low (18-44%), and that cross-state and cross-movie accuracies were the lowest. Results also showed that parcellation resolution and the number of volumes used in each matrix affect fingerprinting accuracies. Next, we calculated three measures of whole-connectome stability for each subject: cross-rest (Rest1-Rest2), cross-state (MovieDM-Rest1), and cross-movie (MovieDM-MovieTP), and three measures of within-state between-subject connectome similarity for Rest1, MovieDM, and MovieTP. We show that stability and similarity were correlated, but that these measures were not related to age. A principal component analysis of these measures yielded two components that we used to test for brain-behavior correlations with IQ, general psychopathology, and social skills measures (n=119). The first component was significantly correlated with the social skills measure (r=-0.26, p=0.005). Post hoc correlations showed that the social skills measure correlated with both cross-rest stability (r=-0.29, p=0.001) and with connectome similarity during MovieDM (r=-0.28, p=0.002). These findings suggest that the stability and similarity of the whole-brain connectome relate to overall brain development, and in particular, to those regions that support social skills. We infer that the development of the functional connectome simultaneously achieves patterns of FC that are distinct at the individual subject level, that are shared across individuals, and that are persistent across states and across runs—features which presumably combine to optimize neural processing during development. Future longitudinal work could reveal the developmental trajectories of stability and similarity of the connectome.

**Highlights:**

- Identification algorithms yielded low accuracies in this developmental sample.
- Individual differences in FC were not as persistent across states or movies.
- Connectome within-subject stability and between-subject similarity were interrelated.
- Stability during rest and similarity during a movie correlate with social skills scores.

## Introduction

### Individually distinct patterns of functional connectivity

Whole-brain functional connectivity (FC), also known as the “functional connectome,” comprises the complete set of correlations amongst BOLD-signal time-courses in the brain. Aspects of FC are highly reproducible at the group level (Damoiseaux et al., 2006; Yeo et al., 2011), but FC matrices also contain patterns that are distinct at the individual subject level (Finn et al., 2015). This has been most clearly demonstrated by studies applying test-retest matching algorithms, which show that an individual subject’s FC matrix is more strongly correlated with another of their own matrices than with an FC matrix from any other subject. Most typically, this is determined by the greatest Pearson’s correlation between pairs of FC matrices, making it possible to identify an individual subject from among a group based only on their connectome (Finn et al., 2017, 2015).

Matching algorithms can be run using data collected under a variety of acquisition states, most commonly resting state and conventional task conditions. In 2017, Vanderwal et al. showed that using movies as an acquisition state for FC measures yielded matching accuracies up to 100% in a small sample of healthy adults; movies preserved and possibly enhanced the identification of individually distinct patterns of FC compared to task-free rest (Vanderwal et al., 2017). A remarkable aspect of the adult human connectome is that FC matrices of individuals can be matched successfully even if those matrices are derived from data collected during markedly different acquisition states. For example, matching across a social action movie (*Ocean’s Eleven)* and a nonverbal movie of abstract shapes and music (*Inscapes)* yielded 94% accuracy, and matching across rest and the action movie achieved 90% accuracy. In a larger sample from the Human Connectome Project, matches across conventional tasks (e.g., working memory task, language task) and rest yielded accuracies of 82-92% (Finn et al., 2015). These results highlight the cross-state persistence of individually distinct patterns in FC in adults.

### Individualization as a maturational process

Recently, this type of identification algorithm was applied to a large pediatric data set (Kaufmann et al., 2017). From a sample of 797 individuals (range 8-22 years, mean 15 ± 3.3), they showed that cross-state identification accuracies across resting state, a working memory task, and an emotion recognition task ranged from 37 to 60%. Using an amalgam measure of identification accuracies across rest and tasks to calculate each participant’s “connectome distinctiveness,” they further showed a robust positive relationship between age and connectome distinctiveness. The authors posited that individualization of the functional connectome is a maturational process that occurs during adolescence, and that this individualization emerges as the connectome becomes stable across conditions.

Connectome stability has typically been investigated as test-retest reliability, and unsurprisingly, pediatric samples have been shown to have lower reliability for multiple whole-brain FC measures relative to adults (Somandepalli et al., 2015; Thomason et al., 2011). Typically, lower reliabilities in pediatric data are thought to reflect variability in data quality or compliance, but here, we build on the findings from Kauffman et al., to ask whether or not connectome stability might be a developmental outcome in and of itself. We hypothesize that within-state (i.e., across different runs of the same condition, such as rest-rest) and cross-state stability are developmental outcomes, and that the lower reliabilities of FC measures in children might, at least in part, be capturing a developmental process rather than simply noise or poor compliance.

### Connectome similarity as a potential developmental measure

Connectome-based identification algorithms rely on the ratio of within- and between-subject correlations. Consequently, we were also interested in asking whether within-state between-subject connectome similarity might be a useful marker of functional brain development. The conceptual developmental model is that if the changes occurring in the developing connectome (e.g., Fair et al., 2008, 2007) are normative changes that optimize overall efficiency, a majority of those pathways and connections might be similar across individuals. Between-subject similarity of the connectome would then be expected to increase as the connectome matures. Here, as an exploratory first step towards examining this idea using cross-sectional data, we investigated the relationship between connectome stability, connectome similarity and behavioral measures of interest.

The present study uses naturalistic conditions to investigate within-subject stability and between-subject similarity of the functional connectome in a psychiatric help-seeking pediatric sample (age range 6-21 years). We used data from two runs of task-free rest and two different movie-watching conditions acquired within a single imaging session from the publicly available *Healthy Brain Network Biobank* (Alexander et al., 2017) to calculate FC matrices for each participant. The matrices were then fed to a test-retest matching algorithm. We had three main predictions. First, we expected to replicate the finding that accuracies for this developmental sample would be lower than those generally obtained for adults across all conditions, especially for cross-state (rest-movie) matches. Second, we hypothesized that secondary whole-brain measures of within-subject connectome stability (cross-rest, cross-movie, and cross-state) would be interrelated, such that an individual participant would be more likely to have higher cross-state stability if they had high cross-rest stability. Third, we predicted that within-subject connectome stability and between-subject connectome similarity would be positively associated with age, and possibly related to behavioral measures of interest. We selected three measures of developmental interest: full-scale IQ, a measure of general psychopathology, and a measure of social skills. We hypothesized that greater stability and similarity would be associated with higher IQ, lower psychopathology scores, and better social skills. In particular, based on the finding that the organization of the pediatric connectome does not persist across states as it does for adults, we thought that cross-state (rest-movie) stability would be most likely to change from childhood to adulthood. We thus hypothesized that cross-state stability would be the most developmentally relevant measure, i.e., that it would relate to behaviors more strongly than the other brain-based measures. Because of the importance of social cognitive development throughout the studied age range, the hubness of “social brain” regions, and the social nature of the movies used, we expected social skills to have the best chance of yielding a significant relationship.

## Methods

### Sample and exclusion criteria

Data from the publicly available Healthy Brain Network (HBN) were used for all analyses (Alexander et al., 2017). This database includes fMRI scans and a battery of cognitive and clinical behavioral scores (as well as data from multiple other modalities including electroencephalography, voice samples, saliva samples, and actigraphy) from a community-based sample from the greater New York area. All participants were seeking psychiatric help. The Chesapeake Institutional Review Board approved all study procedures. Written consent was obtained from all participants 18 years of age and older, and parental consent and participant assent were obtained from all other participants. For this study, data from releases 1-4 were used, and 568 participants were accessed and selected as being from either the Cornell or Rutgers sites, having at least one T1-HCP anatomical scan, as well as data for all four functional runs of interest. Three-hundred and fifty-eight participants were excluded for having a mean framewise displacement of greater than 0.3mm in any functional run, leaving 210 remaining participants (69 females, age range 6-21 years, mean age 12.9 ± 3.6, 128 participants from Rutgers, referred to as HBN-210). The 0.3mm threshold was selected as being mid-range within commonly used thresholds in developmental FC studies (Dosenbach et al., 2017). See Figure 1 for age distribution, head motion statistics, and distribution the Child Behavior Checklist (CBCL) total T scores to reflect the level of psychiatric symptomatology in the sample. A list of subject numbers is available on GitHub at https://github.com/tvanderwal/HBN_stability_similarity.

**Figure 1.**
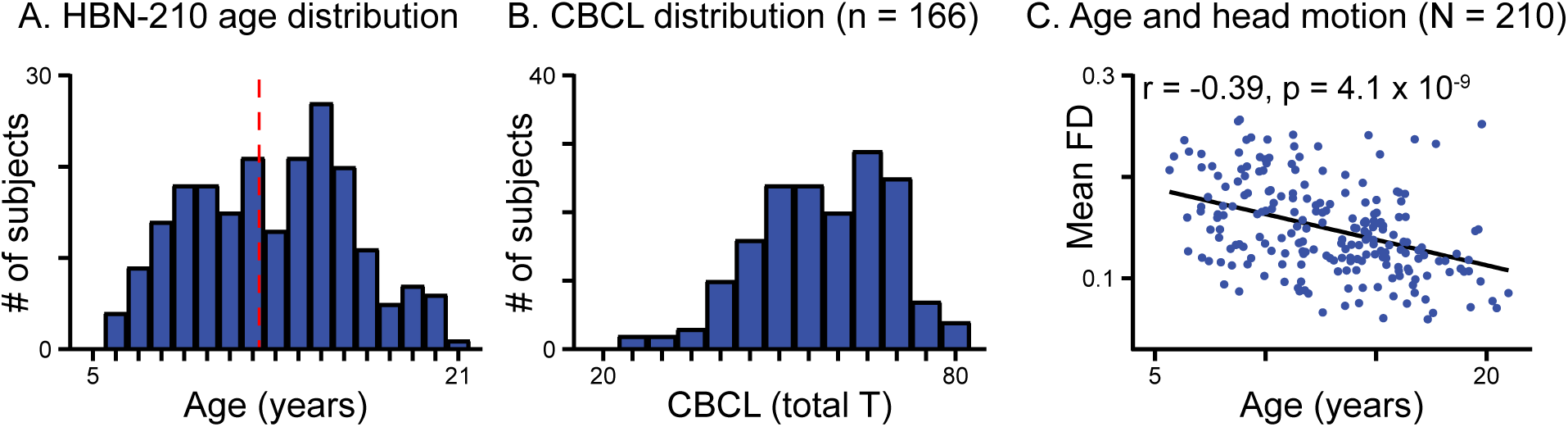
Sample demographics, HBN-210. **A.** Age distribution of the 210 participants (69 females), mean age 12.9 ± 3.6 years (dashed line). **B.** Total T-scores for the Child Behavior Checklist (CBCL), a widely used questionnaire to assess child psychiatric and behavioral problems in which higher scores reflect higher degrees of clinical concern. Scores were available for 166 participants. The right-skew of these data indicates the psychiatric enrichment of the sample, as all participants were seeking psychiatric help (mean total CBCL = 57.2 ± 11.4; clinical cutoff is 65). **C.** As expected, age and decreasing head motion were strongly related. Mean framewise displacement was averaged across the full duration of the four conditions of interest (two resting state runs, and two movie runs) (r = -0.39, p = 4.1 × 10^−9^) for visualization.

### MRI Acquisition

MRI data were collected at Rutgers University using a Siemens 3-Tesla Tim Trio and at the Cornell Brain Imaging Center using a Siemens 3-Tesla Prisma. A 3D T1-weighted structural MP-RAGE volume was acquired (TR=2500ms, TE=3.15ms, FA=8°, slice thickness=0.8mm, slices=224). Echo-planar imaging sequences for the functional runs used TR=800ms, TE=30ms, FA=31°, slice thickness=2.4mm, FOV=204mm, a multiband factor of 6, and voxel size=2.4×2.4×2.4mm.

### Experimental runs

The order of functional runs in the scanning session was fixed, and was as follows: two sequential 5-minute runs of eyes-open rest during which a fixation cross was displayed (Rest1 and Rest2), a 10-minute run during which a clip (with sound) from the movie *Despicable Me* was shown (MovieDM), and a 3.5-minute run during which a short film called *The Present* (also with sound) was shown (MovieTP). The MovieDM clip shows three children without parents asking their begrudging temporary guardian to read them a bedtime story (Coffin and Renaud, 2010). MovieTP is a short animation that tells the story of an unhappy boy who receives a three-legged puppy for a gift (Frey, 2014). Both stories are highly social, feature strong attachment themes, include a mix of poignant and humorous scenes, and were selected to be emotionally provocative. Technical descriptions of the movie clips are available at the HBN study site (http://dx.doi.org/10.15387/CMI_HBN). The first 10 seconds of data were excluded from all conditions, leaving Rest1 and 2 with 375 volumes each, MovieDM with 750 volumes, and MovieTP with 250 volumes. To match the number of volumes across conditions for the main analyses, all functional runs were truncated to 250 volumes. Where indicated, some analyses were conducted with differing numbers of volumes for comparison.

### Phenotypic Assessment

Behavioral measures used in these analyses were collected at visits one and two, with MRI scanning occurring at visit three. Three behavioral measures of interest were selected. To capture global functioning, we used the full-scale intelligence quotient (FSIQ) from the Wechsler Intelligence Scale for Children, Fifth Edition (WISC-V), and the age-adjusted total T-score from the CBCL (Achenbach, 1999). The CBCL consists of 113 items and is filled out by a caregiver. Higher scores indicate greater clinical concern across the full range of psychiatric symptom domains. Third, we selected the total age-adjusted score from the Social Responsiveness Scale-2 (here, SRS) (Constantino et al., 2003). The SRS is a 65-item questionnaire that measures social skills, with higher scores indicating greater impairment. It is classically used in the study of autism spectrum disorders but is also used to capture social problems in other disorders. SRS scores have been shown to be normally distributed in the general population. We focused on a social skills measure because of the highly social nature of the movies used in this experiment, and because of the dynamic social development that is known to occur in the studied age range (Blakemore, 2008; Blakemore and Frith, 2004; Braams and Crone, 2017; Mills et al., 2014). Of the 210 participants, all three behavioral measures were available for 119 participants, 100 of whom were from the Rutgers University site. Demographics for this subset (HBN-119), along with motion statistics and relevant correlations, are shown in Supplementary Figure 1s.

### Data preprocessing

Preprocessing was performed using fMRIPrep version 1.0.15 (Esteban et al., 2019) including slice time correction, motion realignment, and transformation into Montreal Neurological Institute space. Non-aggressive denoising was performed filtering out ICA-AROMA motion comonents, the 6 rigid-body motion correction estimates, framewise displacement, and 6 anatomical CompCor components (Behzadi et al., 2007), and ICA-AROMA motion artifacts. In addition to excluding all subjects with a mean framewise displacement (FD) of greater than 0.3mm in any functional run, ICA-AROMA was used to mitigate motion artifact, as it has been shown to remove spurious noise more effectively than spike regression (Pruim et al., 2015). Motion censoring was not performed to preserve the continuous time-courses and temporal structure of the movie-watching data. Global signal regression was not performed. Data were not spatially smoothed because all planned analyses used a parcellation schema.

### Whole-brain FC matrices

The primary analyses were based on FC connectivity matrices generated using 200 functionally defined regions-of-interest (ROIs) (Craddock et al., 2012). For each subject, the mean time series of each ROI was extracted and Pearson’s correlation coefficients were calculated between all ROI pairs creating a 200×200 whole brain connectivity matrix. Correlation coefficients were then Fisher z-transformed for statistical inference. Duplicate ROI pairs were excluded, leaving 19,900 edges in subsequent analyses. Each subject had an FC matrix for each of the four conditions (Rest1, Rest2, MovieDM and MovieTP).

### Test-retest matching algorithm

The prediction procedure closely followed methods described previously (Finn et al., 2015). Four databases of 210 matrices were created, one per subject for each condition (Rest1, Rest2, MovieDM, MovieTP). The matching algorithm was run between two databases at a time. Each subject’s FC matrix was selected from one database, and the Pearson’s correlation coefficient was calculated between that matrix and every other matrix in the other database. If the matrix with the highest correlation was from that same subject, the pairing was considered a match. Group accuracy of the condition pairing was determined by calculating the proportion of subjects for whom a match was obtained. This analysis was performed across all 12 possible state pairings (e.g., Rest1-Rest2, Rest2-Rest1, MovieDM-Rest1, MovieTP-Rest1, etc.).

To assess the significance of the accuracies produced by the algorithm, we performed nonparametric permutation testing using randomly assigned false identities. The algorithm was run 10,000 times to determine how many times the falsely assigned pair was identified as being the most strongly correlated (Wang et al., 2018). To assess the degree to which head motion might contribute to successful matching, we performed the identification algorithm using only the distribution of framewise displacement values in each run (also from Finn & Shen, 2015). We computed motion distribution vectors for each participant from the framewise displacement time courses of each condition. Sixty bins were set to capture the grand mean of all runs ± 3 standard deviations, and the 1×60 vectors were then used (in place of the FC matrices above) to run the matching algorithm. In this dataset, the grand mean was 0.149mm ± 0.20mm, creating bins between -0.44 and 0.74mm. Limiting this to positive numbers resulted in 38 bins being used. To check for motion effect that might occur due to the small but statistically significant differences in motion across conditions, we created a motion-matched subset of subjects, selecting subjects who had no significant difference in head motion across conditions, and reran the matching algorithm as above using only those subjects’ FC matrices.

To evaluate the effect of condition duration on matching accuracies, we conducted two analyses. First, we maximized the number of available volumes by creating matrices for Rest1, Rest2 and MovieDM for all subjects, this time using the full duration of each condition (instead of truncating at 250 volumes). We also created a new set of matrices by concatenating across the two resting state runs (Rest1+Rest2). The matching algorithm was then rerun with these full-volume matrices and the already full-volume MovieTP matrices. For the second volume-related analysis, we held the amount of data in one matrix constant (Rest1, 375 volumes) and reran the identification algorithm while sequentially increasing the number of volumes in the other three conditions. This process was repeated using the full duration of MovieDM (750 volumes). Here, we wanted to test the assumption that adding volumes to an FC matrix results in linearly increasing matching accuracies.

Finally, to explore the network location of edges that contributed to successful matches, we computed the differential power (DP) of each edge. DP is an empirical probability that indicates the proportion of times a subject is matched to itself rather than to another subject based on that edge. The measure therefore captures how much a particular edge contributes to a successful match (see Finn and Shen 2015 for more details). We then averaged the DP of edges within each network for the Rest1-Rest2, MovieDM-Rest1, and MovieDM-MovieTP pairings. We used the Yeo 7-network parcellation (Yeo et al., 2011), and the network membership of each Crad-200 ROI was assigned to whichever network contained the majority of its voxels. (This Craddock-to-Yeo network key is provided in the paper’s GitHub repository).

### Parcellation resolution

In previous work, we have shown that increasing parcellation resolution can improve identification accuracies, but that it does so somewhat equally across conditions (Vanderwal et al., 2017). To test that finding here, we parcellated the data at all of the 43 resolutions defined in a publicly available atlas (Craddock et al., 2012). The number of clusters ranged from 10-950. We then ran the matching algorithm at each resolution, using the Rest1-Rest2, MovieDM-Rest1, and MovieDM-MovieTP pairings. All other analyses used the Crad-200 parcellation.

### Scan duration

To test the effect of scan duration on the cross-state and cross-run matching accuracies, we ran the identification algorithm while varying the amount of data used, from two to 375 volumes, adding sequential TRs one at a time.

### Number of edges

To test if one condition required fewer edges to facilitate successful identification of individual subjects, we ran the algorithm using an increasing number of edges. Edges were rank-ordered from least contributory (lowest DP) to most contributory (highest DP). We then ran the algorithm using the lowest 0.5% of edges, and successively repeated the procedure adding an additional 0.5% of edges until all edges were included. Arranging edges according to DP rather than spatial location enabled cross-condition comparisons across edges that contributed to accuracy with similar power. If, for example, successful matches could be made with movie-watching data using fewer edges than with resting state data, we would expect a sharp increase in the movie accuracy relative to resting state data at an earlier point on the x-axis.

### Sex-based influence on matching algorithm

To investigate the degree to which having a mixed sex sample influenced the identification accuracies in this developmental sample, we created three cohorts that were matched in sample size, and that did not differ significantly in mean age or head motion. The cohorts were defined using the R MatchIt package for propensity score analysis (Ho et al., 2007). The algorithm was then run separately on each sample following the same procedure used above.

### Connectome stability

We define stability as the edge-wise correlation between a subject’s whole brain FC matrix collected during two separate functional runs. These basic whole-brain correlations were calculated for each subject for cross-run (Rest1-Rest2), cross-movie (MovieDM-MovieTP), and cross-state stability (MovieDM-Rest1). A mean stability measure was also calculated by averaging across these three measures for each subject.

### Connectome similarity

Between-subject similarity is defined as the average of the correlations between a subject’s FC matrix and that of every other subject within a given condition (Rest1, MovieDM, MovieTP). We note that this is an FC-based measure, and therefore can be computed for both movies and rest, as opposed to a time-course based correlation, such as intersubject correlations or ISCs (Hasson et al., 2004).

### Stability-similarity relationships

We then tested for linear relationships between each subject’s stability and similarity measures. Due to the possibility that head motion artifact might still be present in these whole-brain measures of stability and similarity for some participants despite rigorous preprocessing and motion thresholds, we performed follow-up multiple linear regression analyses including each subject’s average framewise displacement from the relevant condition(s) of interest as a covariate.

### Brain-behavior correlations

To test the hypothesis that whole-brain stability and/or similarity is developmentally relevant, we first aimed to reduce the dimensionality of the six measures of connectome stability and similarity by conducting a principal component analysis. The resulting components explaining at least 20% of the variance were selected, and correlations between those components and the three behavioral measures of interest (IQ, CBCL score, and SRS score) were calculated.

## Results

### Head motion

As expected, mean framewise displacement correlated strongly with age (r=-0.39, p=4.1×10^−09^, see Figure 1). Mean FD by condition was Rest1=0.141mm; MovieDM=0.147mm; Rest2=0.15mm; MovieTP=0.16mm). A repeated-measures ANOVA of mean FD across conditions was statistically significant (F(3,209)=6.74, p=0.0002). Differences were driven by greater head motion during MovieTP relative to each of the other conditions (p<0.01). These patterns also held true in the behavioral subsample of 119 participants. Head motion was not correlated with any of the three behavioral measures of interest (see Supplementary Figure 1s).

### Accuracies of identification algorithm

For matches across all conditions, matching accuracies ranged from 18-44% (Figure 2A), whereas permutation testing using falsely assigned subject pairs yielded accuracies of 0.3-0.4% (Supplementary Figure 2s). The only pairing of a truly repeated condition (Rest1 and Rest2) had the highest accuracies (43% and 44%), while the four cross-state (Movie-Rest) pairings had the lowest (18-36%). The two cross-movie accuracies were comparable (27% and 34%). To statistically assess these differences in accuracy, we performed permutation testing. Condition labels were randomly shuffled for each subject in each permutation. Accuracy was calculated for all condition pairs (each one a random shuffling of actual conditions) and subsequent pair-wise differences were computed across the accuracy matrix. The resulting 60,000 differences (or deltas) were ordered from smallest to greatest to form the null distribution. The top 2.5^th^ percentile of these deltas represented a p<0.05 (two-tailed) likelihood that a given pair of accuracies were different, which corresponded to a difference in accuracies of 8.6% and above.

**Figure 2.**
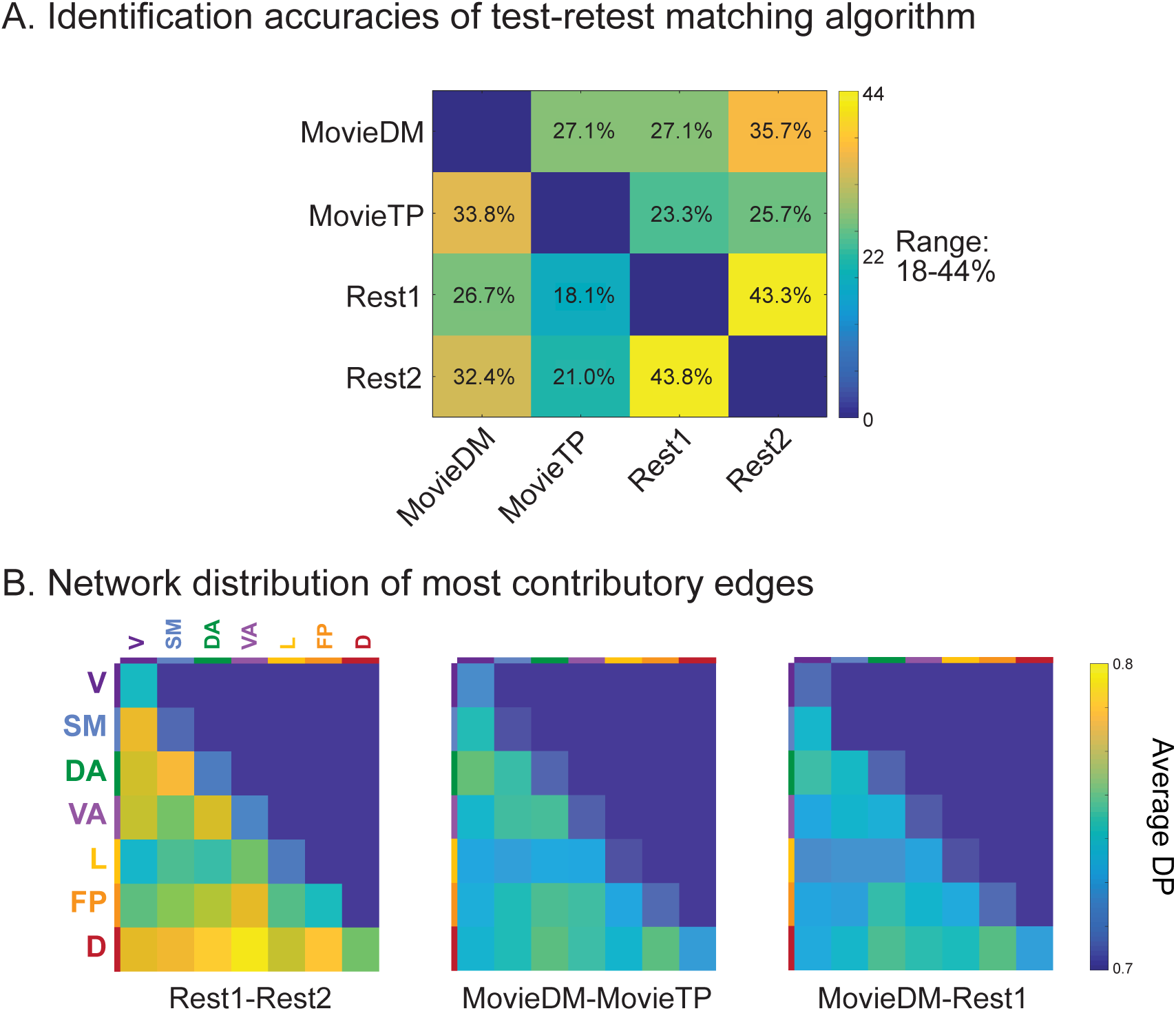
Test-retest matching accuracies, HBN-210. **A)** Matching accuracies of the identification algorithm ranged from 18-44%, with the highest accuracies attained by the only true condition pairing, Rest1-Rest2 (and vice versa). Movie-TP pairings yielded the lowest accuracies (18-23%), and cross-movie and MovieDM-Rest pairings were in the middle (26-36%). Permutation testing showed that differences of greater than 8.7% correspond to p<0.05. **B)** When examining the distribution of the contributory edges by looking at the mean differential power (DP) by network, between-network edges contribute to successful identifications more often than within-network edges. As has been previously shown, heteromodal cortex such as the frontoparietal, default and dorsal attention networks exhibited high mean DP, but overall, DP appears to be widely distributed. Condition-based differences appear to be a matter of degree rather than demonstrating clear condition-by-network effects.

When the algorithm was re-run using the full duration of each scan to compute FC matrices for each condition (i.e., not limited by the 250 volumes of MovieTP), the accuracies shifted, with a new range of 20-49%. In most cases, pairings of mismatched durations got worse (e.g., MovieDM-MovieTP went from 27 to 20%), and pairings with longer durations improved (e.g. Rest1-Rest2 accuracies went from 43 to 48%). Via permutation testing, the only shifts that reached significance were decreases in accuracy for both of the cross-movie pairings. Because of these decreases, we performed a second analysis where the number of volumes in one matrix was held constant, and the algorithm was repeatedly rerun as volumes were sequentially added to the other matrix. This showed that adding data does not result in a linear increase in accuracies in all cases, and in fact, some trends appear to be nonlinear. When using motion distribution parameters alone to match subjects, accuracies ranged from 2-8%, and when using motion-matched groups (n=190), percentages ranged from 20-46% and followed the same patterns observed in the full sample. Results of these supporting analyses are shown in Supplementary Figure 2s.

When assessing the network distribution of DP, we found that between-network edges contributed to successful identification more often than within-network edges. As has been previously shown, heteromodal cortex such as the frontoparietal, default and dorsal attention networks exhibited high mean DP (Finn et al., 2015; Vanderwal et al., 2017), which may in part be due to high reliability of FC in these regions (O’Connor et al., 2017). Overall, DP appears to be widely distributed. Condition-based differences appear to be a matter of degree rather than demonstrating clear condition-by-network effects.

### Parcellation resolution

As expected, when testing with different parcellation resolutions, identification accuracies increased as parcellation resolution increased for all pairings (cross-run, cross movie, and cross-state) (see Figure 3A). The highest accuracy attained was for Rest1-Rest2 pairings, which was 60%. All other analyses presented here used the Crad-200 parcellation.

**Figure 3.**
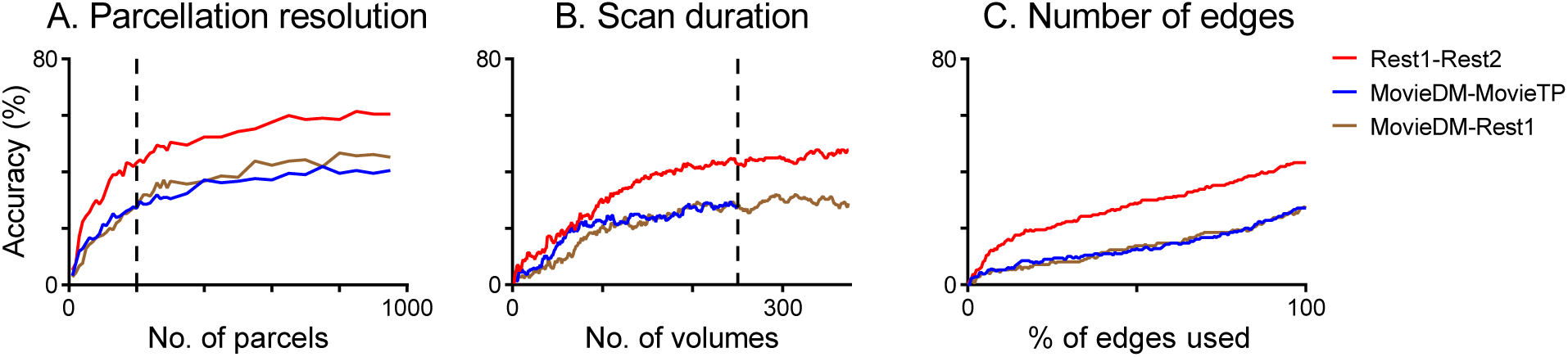
Effects of processing parameters on identification accuracies across different pairings. Altering preprocessing parameters did not change the overall pattern of results. The only properly matched pairing, Rest1-Rest2 (red line), achieved higher accuracies than either the cross-movie or the cross-state pairings, regardless of the parcellation resolution, the number of volumes, or the number of edges used. The dashed line in A denotes the Crad-200 parcellation, and in B denotes the truncated 250 volumes, both of which were used for all other analyses.

### Scan duration

Within the range of available volumes, longer scan durations improved matching accuracies (Figure 3B). After 100 volumes, accuracies appear to increase incrementally with added volumes, and the relationships across conditions (i.e., cross-rest accuracies are greater than both cross-movie and cross-state matches, which are similar to each other) appear consistent. We also point to Supplementary Figure 3E for different results when volumes beyond the studied range were included.

### Number of edges used

Including more edges yielded higher accuracies overall. Again, the trajectory for the Rest1-Rest2 pairing separates out from both the cross-state and cross-movie pairings (Figure 3C).

### Sex-based influence on matching algorithm

The algorithm was rerun using three separate cohorts (females, males, mixed, all n=69). As expected, matching accuracies were higher than in the original analysis (N=210), but this was true and similar across all three cohorts, so was attributed to sample size rather than to sex-based differences.

To statistically assess the differences in accuracy across the three cohorts, we again performed permutation testing, this time extracting two random groups of 69 participants without replacement from HBN-210. For each iteration of the 1000 iterations, we reran the identification algorithm separately on both random cohorts of 69 participants, and determined the differences (or deltas) in accuracy within each condition across the cohorts. Null distributions of these differences were created for each condition pair, where the top 2.5th percentile represented p<0.05 (two tailed). The smallest possible difference across all distributions that reached significance was 13%, while the largest difference in the three actual cohorts was 11.6%. Matching accuracies were therefore judged to be comparable across the cohorts, indicating that using a mixed sex sample for the main analysis did not substantially impact the results even in this developmental sample. The cross-condition pattern of matching (i.e., the overall color layout on the matrices) was also comparable to the main analysis. These results are shown in Supplementary Figure 3s.

### Connectome stability and similarity

Figure 4 illustrates how stability and similarity measures were calculated, and how they relate. As expected, all six measures were significantly correlated with head motion (see Supplementary Figure 4s), so to investigate relationships among these measures, we ran a linear regression model with motion as a covariate. The motion covariate was created by averaging the mean FD across all runs that were used in computing the resulting brain metric (e.g., for cross-rest stability, the mean FDs from Rest1 and Rest2 were averaged). The partial correlation coefficient and accompanying p-values are reported for the variables of interest. We found that all pairwise measures of stability were positively correlated as follows: cross-rest and cross-state, r=0.62, p=9.2×10^−24^; cross-state and cross movie, r=0.62, p=9.2×10^−24^; cross-rest and cross-movie, r=0.32, p=1.6×10^−6^). Between-subject similarity during Rest1 and MovieDM were also correlated (r=0.34, p=6.0×10^−7^). When each subject’s three stability scores and their three similarity scores were averaged, mean stability and mean similarity were also strongly correlated (r=0.65, p=6.5×10^−26^), showing that participants with higher within-subject stability also demonstrated higher between-subject similarity. Contrary to predictions, no significant relationships with age were found when testing for either linear or quadratic fits (see Supplementary Figure 5s). Further, neither mean stability nor mean similarity were correlated with age (stability: r=-0.0001, p=0.99; similarity: r=0.0106, p=0.91). For completeness, all pair-wise correlations among the six secondary brain measures with each other without covarying for head motion—and correlations with head motion and age—are provided in Supplementary Figure 4s.

**Figure 4.**
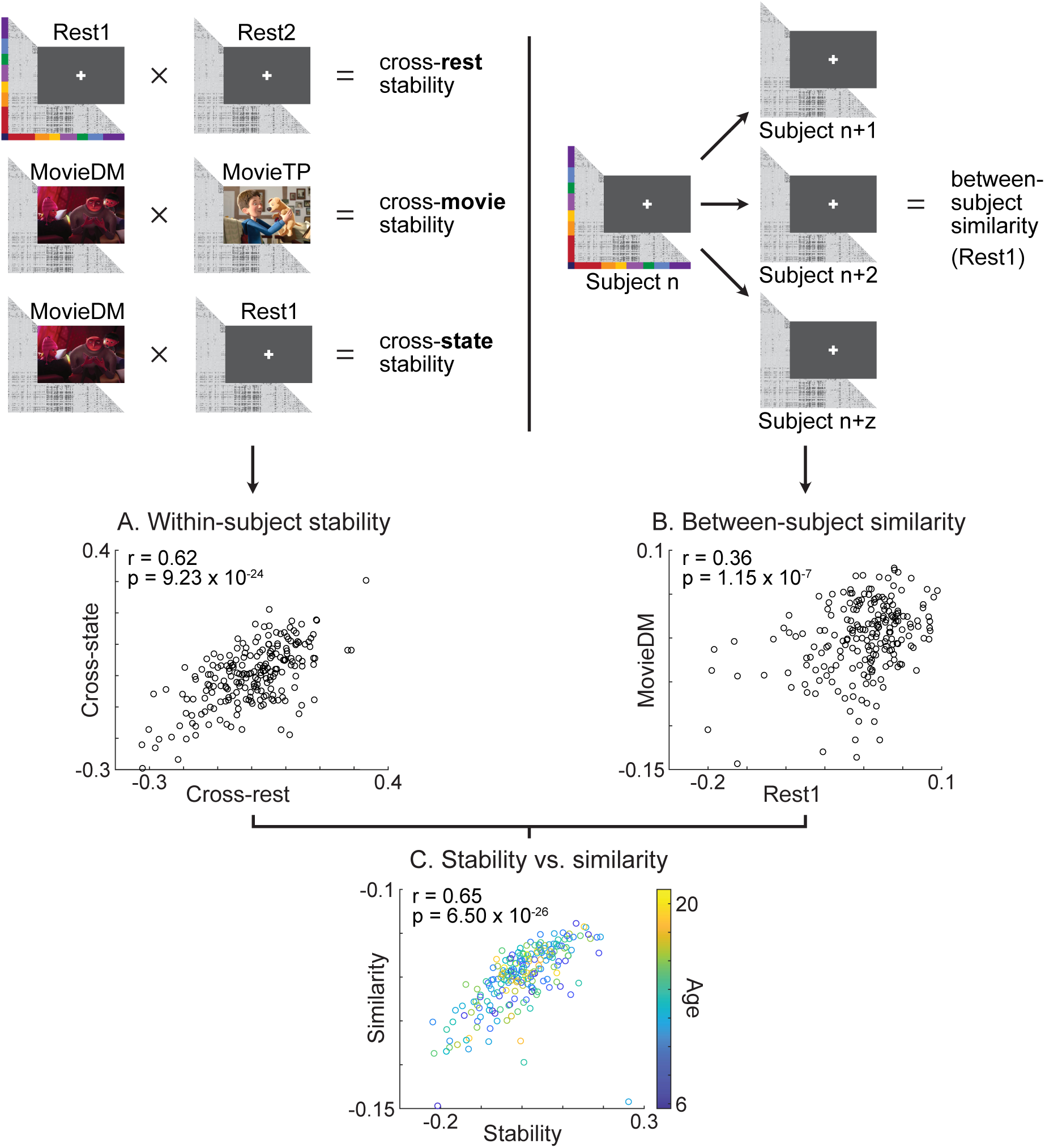
Whole-brain within- and between-subject measures of FC correlations, N=210. Six measures of connectome stability and similarity were calculated for each subject. Plot A shows the relationship of cross-rest and cross-state within-subject connectome stability. For between-subject similarity, we calculated the correlation between each subject’s matrix and a matrix from every other subject for each condition. Each subject’s similarity value is the average of those between-subject correlations for a condition. Plot B shows the relationship of between-subject connectome similarity for MovieDM and Rest1. For C, each subjects’ three measures of stability and similarity were averaged and plotted, with age mapped onto that relationship using color. All measures plotted are the residuals after regression of head motion. Correlations without covariates between all six measures, and with head motion and age, are shown in supplementary Figure 4s.

We were still concerned that the apparent interrelated nature of these measures could be driven by head motion, such that a participant with low stability due to noise from head motion would also be expected to be less similar to the other participants because of that same noisiness. As a second test, we binned participants by mean FD, and selected a range of bins to get the most subjects with the least spread in FD. Using FD of 0.13-0.15mm, we had 33 participants. Neither stability nor similarity in this subsample was significantly correlated with FD, but the robust correlation between stability and similarity persisted (r=0.81, p=1.17×10^−8^).

### Brain-Behavior Correlations

Due to the interrelatedness of the six measures of connectome stability and similarity, a principal component analysis (PCA) was used to reduce dimensionality. The PCA yielded components that explained 3-60% of the variance. The top two components explained 59% and 24% of the variance and were selected for subsequent analyses (see Figure 5 for scree plot and component loadings). Component 1 was weighted toward the within-subject stability measures, and Component 2 was weighted towards the movie-related measures, particularly cross-movie stability. Age did not significantly predict either component in a linear regression model, also using sex, site, and head motion (Component 1, partial r=-0.091, p=0.33; Component 2, partial r= 0.14, p=0.15).

**Figure 5.**
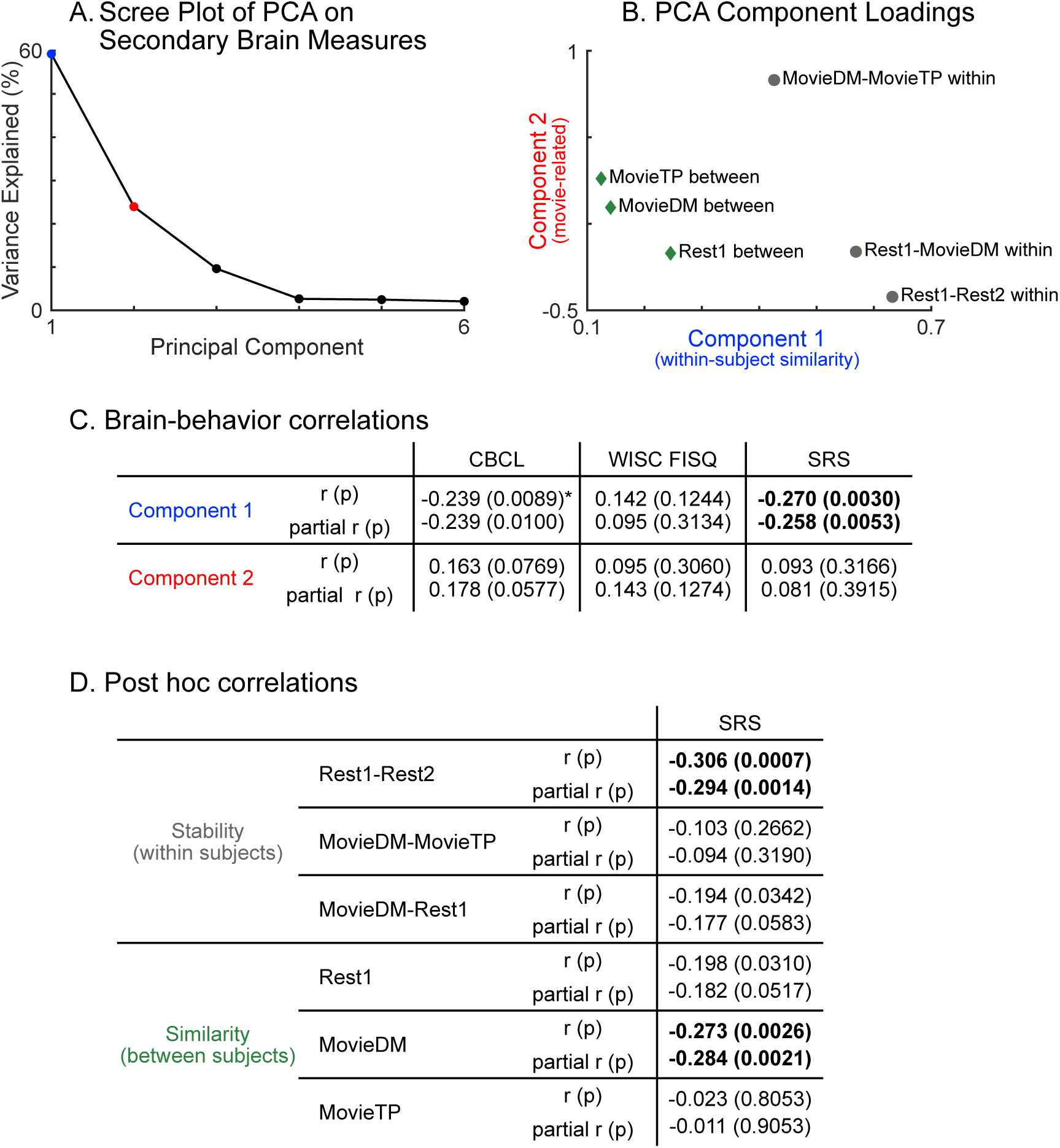
Brain-Behavior Correlations, n=119. **5A)** A Principal Component Analysis performed on the six secondary brain measures (from Figure 4) yielded the scree plot shown here. The top two components were selected for further analyses**. 5B)** Component 1 explained 59% of the variance and was weighted towards the within-subject stability measures. Component 2 explained 24% of the variance, and was weighted towards movie-related measures, especially cross-movie stability. We then tested the two components for correlations with the three developmental measures of interest, CBCL (general psychopathology and behavioral problems), full-scale IQ, and a social responsiveness score (SRS). We report standard correlations on rows denoted “r/p”, and partial correlations after including head motion, age, sex and site as covariates, on rows denoted “partial r/p.” **5C)** A significant relationship was identified between component 1 (FC stability) and social skills. We also note the strong relationship (with asterisk) between CBCL and Component 1 that did not reach statistical significance after Bonferroni correction. **5D)** Post-hoc statistical tests between SRS and the 6 secondary brain measures showed significant relationships between SRS and cross-rest stability, and between SRS and MovieDM similarity (Bonferroni corrected, p<0.008). Directionality of the brain-behavior correlations indicate that higher stability or similarity are associated with lower SRS scores (better social skills).

To investigate relationships between these components and behavior, we ran linear regression models predicting each component from the behavioral measures of interest. Sex, site, and motion covariates were also included. Partial correlation coefficients and accompanying p-values are reported for the regressors of interest. When using age-adjusted measures of psychopathology and general behavioral problems (CBCL), full-scale IQ (FSIQ), and social responsiveness score (SRS), Component 1 was significantly correlated with SRS (r=-0.26, p=0.005). Though it did not meet significance with Bonferroni correction, we also note a strong correlation between Component 1 and CBCL (r=-0.24, p=0.01) (Figure 5C). None of the other relationships between brain components and behavior were significant. CBCL and SRS were subsequently found to be strongly correlated (r=0.71, p=3.8×10^−19^) and all CBCL subsets (e.g., Social Problems, Attention Problems, Somatic Complaints, etc.) were significantly correlated with SRS. Applying a median split to the sample, as well as splitting the CBCL into high and low clinical concern subcohorts (based on the cutoff score of 65), showed that the correlations between CBCL and SRS were still significant in all splits, and therefore not skewed by those participants with more severe symptomatology. When the six stability and similarity measures were correlated with either CBCL or SRS while controlling for the other, no significant relationships were observed. The association between CBCL and SRS has been shown previously, especially in clinical samples (Constantino and Frazier, 2013; Hus et al., 2013). Because of this, and the subthreshold significance of the component-CBCL correlation, SRS was the only variable carried forward for post hoc tests.

We then conducted post hoc multiple linear regressions to test for relationships between SRS and the six brain measures of stability and similarity. Again we report both the correlations (r- and p-values) of these pair-wise correlations, as well as the partial r- and p-values from a linear model that included head motion, age, sex, and site as covariates (Figure 5D). We found that within-subject stability for rest was significantly correlated with SRS (r=-0.29, p=0.001). Between-subject similarity during MovieDM was also significantly correlated with SRS (r=-0.28, p=0.002).

## Discussion

This study investigated within-subject stability and between-subject similarity of the whole-brain connectome during movies and rest in a psychiatrically enriched sample of 210 children and adolescents, ages 6-21 years. As expected, we replicated the finding that matching accuracies for this developmental sample were low (Kaufmann et al., 2017). We also extended the finding of low cross-state stability in children to include naturalistic conditions, showing that both cross-movie and cross-state (movie-rest) matching accuracies were particularly low in this sample. Also as hypothesized, we showed that within-subject connectome stability (cross-rest, cross-movie, and cross-state) were correlated with each other, meaning that an individual participant was more likely to have higher cross-state stability if they had high cross-rest stability. The same was also true for between-subject connectome similarity. Additionally, connectome stability and similarity were shown to be largely interrelated measures, even when accounting for head motion. Contrary to our hypotheses, we found no relationship between either connectome stability or connectome similarity and age. In a subset of 119 participants, we identified significant brain-behavior relationships using the stability and similarity measures: cross-rest stability was significantly correlated with the SRS, an index of social skills. Further, connectome similarity during the movie *Despicable Me* was significantly correlated with social skills. Overall, we conclude that stability and similarity of the functional connectome may be meaningful developmental markers, but multiple methodological questions and caveats remain that can be especially influential when conducting these types of analyses in children and adolescents.

### Fingerprinting in developmental samples

We showed that test-retest matching algorithms of whole-brain functional connectivity matrices in a pediatric sample are low overall, and that accuracies for cross-state and cross-movie matches are especially low. For example, matching accuracies across two different cartoons from the same scan session in this study were 27% and 34%. In a previous study in healthy adults (albeit with a smaller sample of 31 participants), we found cross-movie matching accuracies to be between 80-94%, even across markedly different movies and with one-week interscan intervals (Vanderwal et al., 2017). In other words, patterns of functional connectivity (FC) that are distinct at the individual subject level appear to persist (or are identifiable) across states for healthy adults, but much less so in this developmental, psychiatrically enriched sample. This observation supports the idea that connectome stability and similarity are different in children than adults. Additionally, these data show that applying fingerprinting algorithms in developmental and/or psychiatric samples raise methodological considerations. Here we discuss six relevant issues.

A first potential caveat to this finding relates to scan duration, or the number of volumes used to construct each matrix. The main analyses used matrices that were all truncated at 250 volumes to match the shortest movie clip (MovieTP). When we re-ran the matching algorithm using the full duration of available volumes for Rest1, Rest2 (375 volumes), and MovieDM (750 volumes), accuracies shifted both up and down, and the only shifts judged to be significant were decreases in accuracy. Furthermore, whereas we had hypothesized that adding data would result in an increase in accuracies across the board, incrementally changing the number of volumes in one matrix of a paired condition shifted accuracies nonlinearly in some cases (Supplementary Figure 2s, panel E). Together, these results indicate that the identification of individually distinct patterns of FC depends on the amount of data used to construct the FC matrices and that more data does not necessarily confer an increase in fingerprinting accuracies. It would be useful in future work to determine whether there are particular thresholds for data quantity for fingerprinting algorithms, and whether these differ in children versus adults, and across states (or even different movies). Given the limitations of the data used here, it is possible that individually distinct patterns of FC are present in children to the same degree as in adults, but that more data would be required to identify them in children.

A second finding was that increasing the parcellation resolution increased identification accuracies for all combinations of conditions (Figure 3B). For example, the rest-rest pairing increased from 40% at 250 clusters to 60% at 950 clusters, which suggests that parcellation resolution is a nontrivial issue. This may relate to improved reliability of FC at higher parcellations (Xu et al., 2016). Here, we used an adult-based resting-state parcellation scheme (Craddock et al., 2012). In a developmental sample, this could be a significant issue, and using better parcellations that are defined within the study sample (Shen et al., 2013), or that are condition-specific (Salehi et al., 2019) may reveal that individually distinct patterns in FC are robustly present during childhood if small differences in topography are better accounted for. Another potential avenue to fully optimize this approach would be to use hyperalignment (Conroy et al., 2013; Guntupalli et al., 2018; Haxby et al., 2011; Nastase et al., 2019) in a developmental sample, though the data duration (upwards of 20 minutes) required for this approach—even with movie-watching— remains a challenge.

A third limitation was that the only true matched pairing available for the algorithm was between Rest1 and Rest2, as the HBN data set does not include two runs of the same movie. One of the hypotheses that originally motivated this study was that individually distinct patterns of functional connectivity may exist in children when the brain is being driven under naturalistic conditions, but that those connectivity patterns do not yet (either via Hebbian learning or repeated use, etc. (Keysers and Perrett, 2004)) persist as part of the intrinsic functional connectome. Whether the pediatric connectome is markedly more stable across runs of the same movie relative to rest, or relatedly, whether matching accuracies would be higher for pairings using the same movie in children, remain open questions.

Fourth, head motion artifact is a major issue in any developmental study of functional connectivity (Grayson and Fair, 2017; Power et al., 2012), and because fingerprinting algorithms rely on a ratio of within and between subject measures of FC, head motion may be confounding in complex ways. The head motion treatment in these analyses was quite strict: we excluded any participant with greater than 0.3mm mean framewise displacement, even if they only exceeded the threshold in a single run. We also used ICA-AROMA to remove motion-related artifact (Pruim et al., 2015), tested for residual motion effects by running the algorithm with just motion-based outputs (Finn et al., 2015), and in a subsample where motion across conditions was matched. However, as below, cross-run and cross-state stability were correlated with motion, and insofar as these measures are the building blocks of a fingerprinting algorithm, and due to the robust and intractable relationship between age and head motion, the influence of head motion on these results remains a concern. Future studies in high-motion groups that use movie-watching could help determine whether motion correction strategies that leave the time-course intact (as opposed to motion censoring in which volumes are removed) are superior and could further investigate the role of motion artifact in these types of algorithms.

Fifth, low accuracies could be due to issues with movie choices and, in some cases, the order within the scanning session. *The Present* was shown last during the scanning session, and it is possible that the data for MovieTP suffered because of this order effect even beyond the movie’s elevated head motion. Additionally, it is possible that idiosyncratic patterns of FC were evoked by these particular movies, and though our previous work in adults suggests this should not be the case, additional work is needed to rule out movie-specific effects on fingerprinting algorithms.

Finally, though we tested for the effects of some preprocessing parameters (Figure 3), other parameters, such as the use of global signal regression, remain untested. While some fingerprinting papers used GSR (Finn et al., 2015), others (including this one) did not (Kaufmann et al., 2017; Vanderwal et al., 2017), which potentially confounds comparisons of results.

### Connectome stability and similarity as developmental measures

We calculated six secondary measures of whole-brain correlations: three within-subject measures of FC stability (cross-rest, cross-state, and cross-movie), and three between-subject measures of FC similarity during Rest1, MovieDM, and MovieTP. Analysis of these measures revealed that the different types of stability are correlated, as are similarity under different conditions. Further, if a participant had high within-subject stability, they generally also exhibited a higher degree of similarity with the connectomes of the other participants (Figure 4C). These results suggest that these organizational characteristics are interrelated, and possibly interdependent, in the maturation of the pediatric connectome. Put differently, we speculate that many of the patterns of FC that emerge during development (Fair et al., 2007; Grayson and Fair, 2017) do so to optimize increasingly complex information processing, and that because the processes being optimized are largely shared, many optimized pathways are similar across individuals. When a connectome reaches that “optimized” set of patterns (through repetition or FC reorganization, however that might happen), that connectome would also be more stable or persistent across time and across states. Within this conceptual framework, neural maturity of FC relationships in a subset of edges would be reflected in both stability and similarity.

A remaining consideration is that some form of noise or diminished signal reliability could explain the interrelatedness of these measures. Both stability and similarity were significantly correlated with head motion. However, in a linear model, when head motion was covaried, the relationship remained significant. We also selected a cohort of subjects with minimal range of FD (0.13-0.15mm, n=33). In this subcohort, FD was not correlated with either stability or similarity, but stability and similarity remained strongly correlated. None of these analyses are definitive, however, and questions about the role of head motion in our findings remain. We cannot exclude other confounds, including attentional differences, varying levels of arousal or actual sleep; such confounds may be more problematic than usual in developmental, psychiatric samples.

Aspects of neural stability, or conversely, within-subject measures of variability, have long been of interest in developmental research. McIntosh et al. argue that neural variability provides options and efficiency to the brain. They showed that variability of evoked potentials during task-based EEG increased with age, and that increasing neural variability accompanies decreasing behavioral variability and improved task performance (McIntosh et al., 2008). Similarly, using fMRI, Hutchison et al. investigated the dynamics of FC organization during task and rest in participants ages 9-30 years old (Hutchison and Morton, 2015). During rest, adults and children expressed the same number of network configurations or states, but during task, adults exhibited a larger repertoire of states, as well as more (and faster) transitions between states. However, in participants across ages 9-32 years, Marusak et al. found a positive association between age and the temporal variability between functional connectivity networks during rest (Marusak et al., 2017). In an early pediatric movie-watching studies, Emerson et al. compared FC during rest and movie-watching in healthy six-year old children (Emerson et al., 2015). Using both independent component and seed-based analyses, they showed that three networks (frontal control, default and dorsal attention) reconfigured during movie-watching such that frontal control-default network connectivity was upregulated, and frontal control to dorsal attention network connectivity was downregulated. They observed the same pattern in adults but noted that the amount of change from resting state to movie-watching (i.e., the cross-state shift in FC) was less pronounced for children relative to adults. While these findings do not support the idea that cross-state differences are more pronounced for children, they do suggest that age-based differences in FC organization are more prominent during task/movie than resting state in typical development. They also underscore the importance of investigating cross-state differences in FC across development.

### Social scores and functional connectivity

Using a principal component analysis, we showed that the variance captured in these large-scale, interrelated secondary brain measures of stability and similarity relates to a broad measure of social skills, the SRS-2. Post hoc analyses showed that better social skills correlated with greater stability across resting state runs (r=-0.029, p=0.001), and also with greater between-subject similarity of the connectome during the *Despicable Me* movie run (r=-.028, p=0.002). We did not find relationships with age or IQ, and we did not find significant relationships with the other movie data (*The Present*).

The finding that two of the whole-brain secondary measures of stability and similarity correlated with SRS scores meshes with the central role that social cognition plays in the developing brain. We had hypothesized that stability and similarity might related to chronological development, in accordance with the overarching idea that as FC relationships become more efficient and mature, they become more stable within a person and more reflective of patterns that are shared across people. With regards to social skills, marked improvement in social cognitive skills during childhood and adolescence is generally considered to be the dominant developmental focus of this highly dynamic age range (Blakemore, 2008). The brain regions implicated in “the social brain” are among the most densely connected in the cortex (McCormick et al., 2018; Oldham and Fornito, 2019; Saxe et al., 2009), and previous studies have shown that significant changes in FC organization of these regions occur during development (Richardson et al., 2018). Recently, Lake et al. used Connectome Predictive Modeling to identify a complex set of FC edges that predicted SRS scores in a large sample of children and adolescents with and without autism spectrum disorder (Lake et al., 2019). The SRS network identified by the machine-learning algorithm was predictive of SRS scores of unseen participants, even when those participants were from a separate study and had attention-deficit hyperactivity disorder. These transdiagnostic results indicate a surprisingly tight coupling between intrinsic functional connectivity and social skills. Our findings here support the idea that SRS scores relate closely to large-scale functional connectivity relationships in the developing brain, both during rest and movie-watching conditions.

One complication of the brain-behavior correlations we identified is that, contrary to our hypothesis, stability and similarity were not correlated with age. Moreover, none of the behavioral measures tested were correlated with age, and the only robust relationship observed with age was head motion. The brain-behavior correlation analyses were conducted with only 119 participants, and given the clinical heterogeneity of the sample, it is possible that age-relationships would emerge with a larger sample. It is also possible that because the sample is psychiatrically enriched, differences in neurological maturation may have made chronological age less meaningful as a developmental marker. That is admittedly a controversial suggestion, but it implies that measures such as SRS might be more developmentally meaningful than age. Other possible contributing limitations that we cannot exclude include scan duration, movie choice, choice of motion criteria, the lack of two runs of the same movie, and possibly the lack of global signal regression (Li et al., 2019). Either way, further studies with larger samples are needed to test the brain-behavior relationship we observed.

We also note that the between-subject similarity during movies were correlated with SRS scores only during one of the movies. We note differences with the stimuli as possible explanations for this finding. As above, *The Present* came last in the scanning session, and there are also relevant differences between the movies themselves. For example, *Despicable Me* is a social, highly verbal movie that features children of different ages and sexes. *The Present* is largely nonverbal and features a single young adolescent male. *Despicable Me* may have been more engaging and more likely to drive the “whole-brain” overall, and this combined with the order effect of MovieTP, may have made the FC data derived from *Despicable Me* more meaningful. A more sobering possibility is that the SRS relationship might stem from some idiosyncratic, stimulus-specific aspect of MovieDM (Westfall et al., 2017), though this concern would not apply to the cross-rest stability correlation. Overall, these findings suggest that the developmental changes in neural circuitry that produce cross-rest stability and between-subject similarity of the functional connectome may also be relevant for supporting the development of social skills, arguably the most complex processing which the brain performs.

### Conclusions

Based on the fingerprinting findings here, and on the subsequent analyses using whole-brain measures of stability and similarity, we suggest that the developmental reorganization of the functional connectome during childhood and adolescence is accomplishing congruent trajectories, meaning it is becoming both more stable within-subjects (across resting state runs and across states or conditions) and more similar between-subjects in a potentially interrelated way. Moreover, it seems that a connectome that is both stable and similar also contains individually distinct patterns of FC—perhaps as a byproduct of attaining the rich neural variability that maximizes efficiency within an individual’s brain. The strongest finding in these data was that the stability of the connectome across resting state runs relates to a measure of social skills, and we speculate that the same processes that support the stabilization of the connectome also support social cognitive development. Nevertheless, significant caveats attend these findings. Particularly because this sample was both developmental and psychiatrically enriched, and because these findings did not relate to age, further investigation is needed. These data also underscore the importance of both the parcellation resolution and the number of volumes used in each matrix in fingerprinting algorithms and highlight the need for large-scale datasets with multiple, long movie runs. Ideally, longitudinal data would be used to characterize the trajectory of connectome stability both across rest runs and across different states—and to track how stability relates to both connectome similarity and behavioral measures of development—over time.

---

**Figure 1s:**
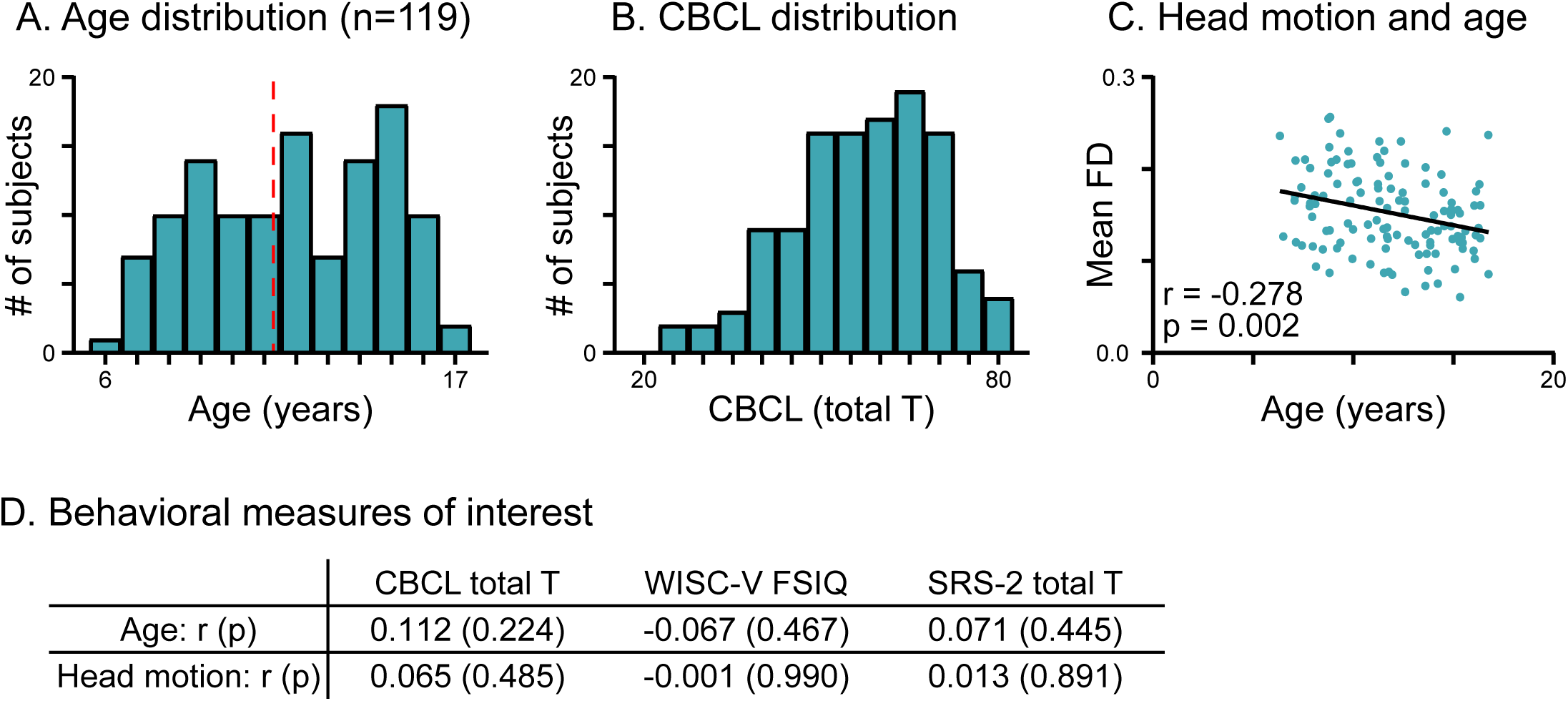
Demographics of behavioral subsample (n=119)

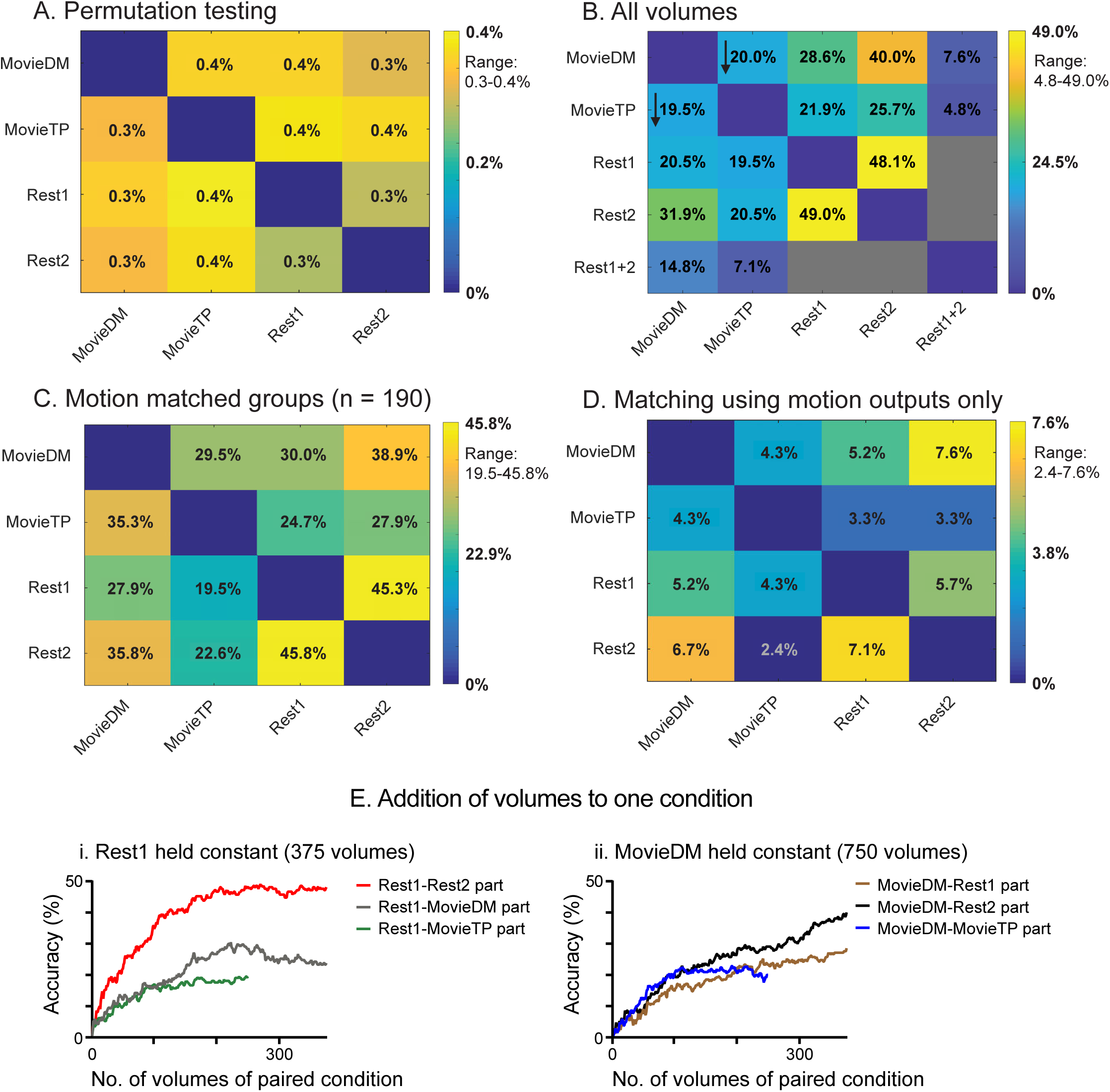

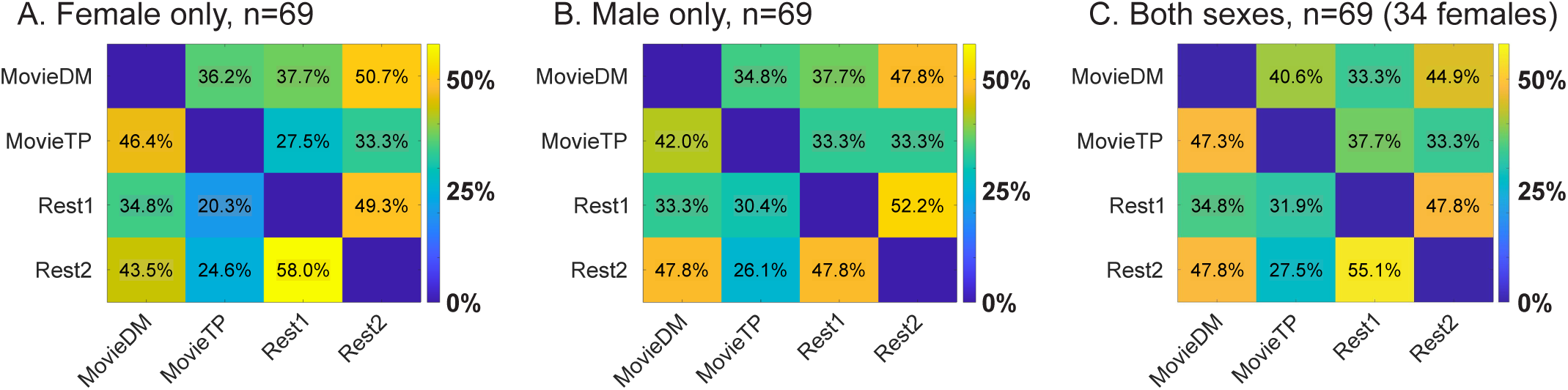

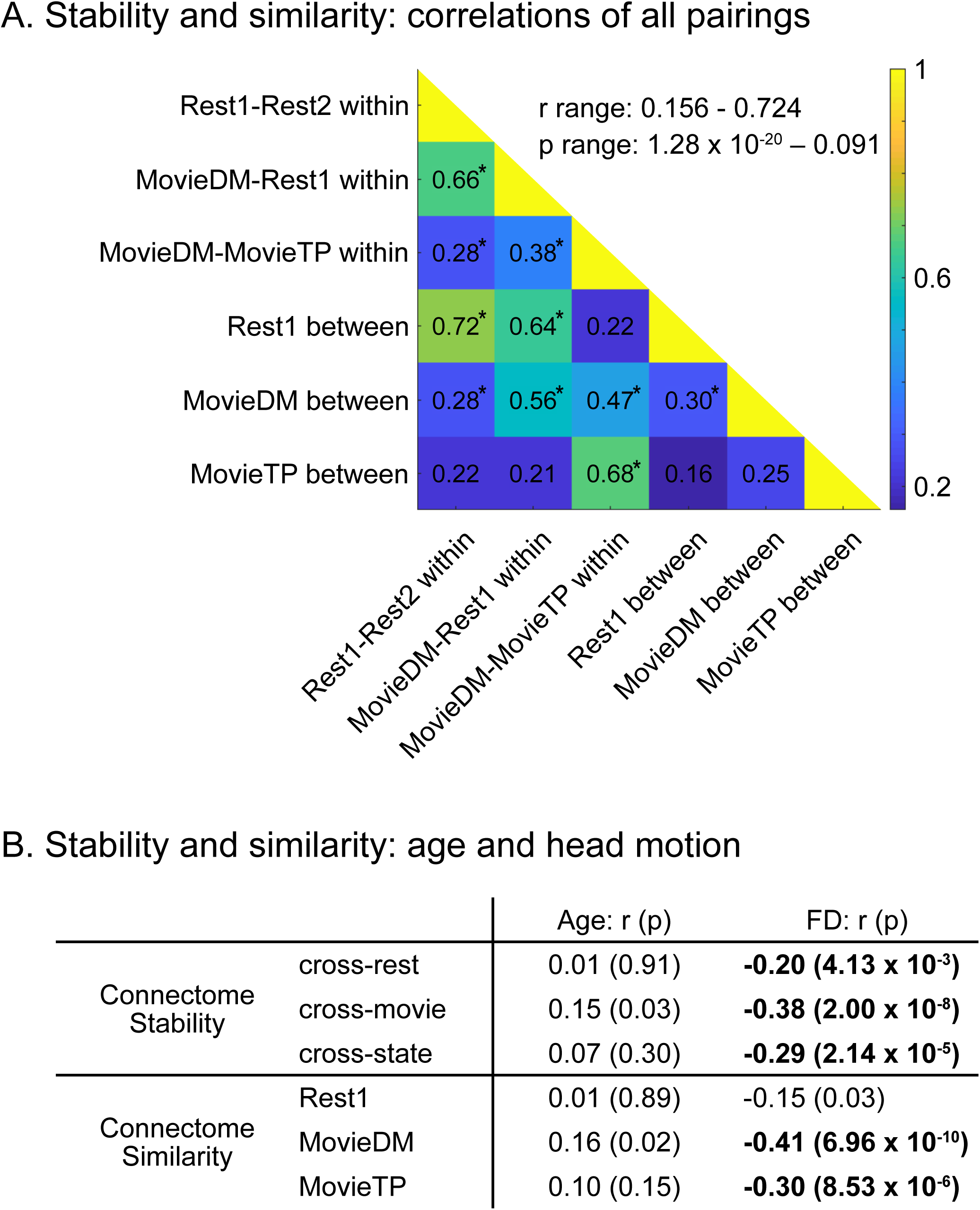

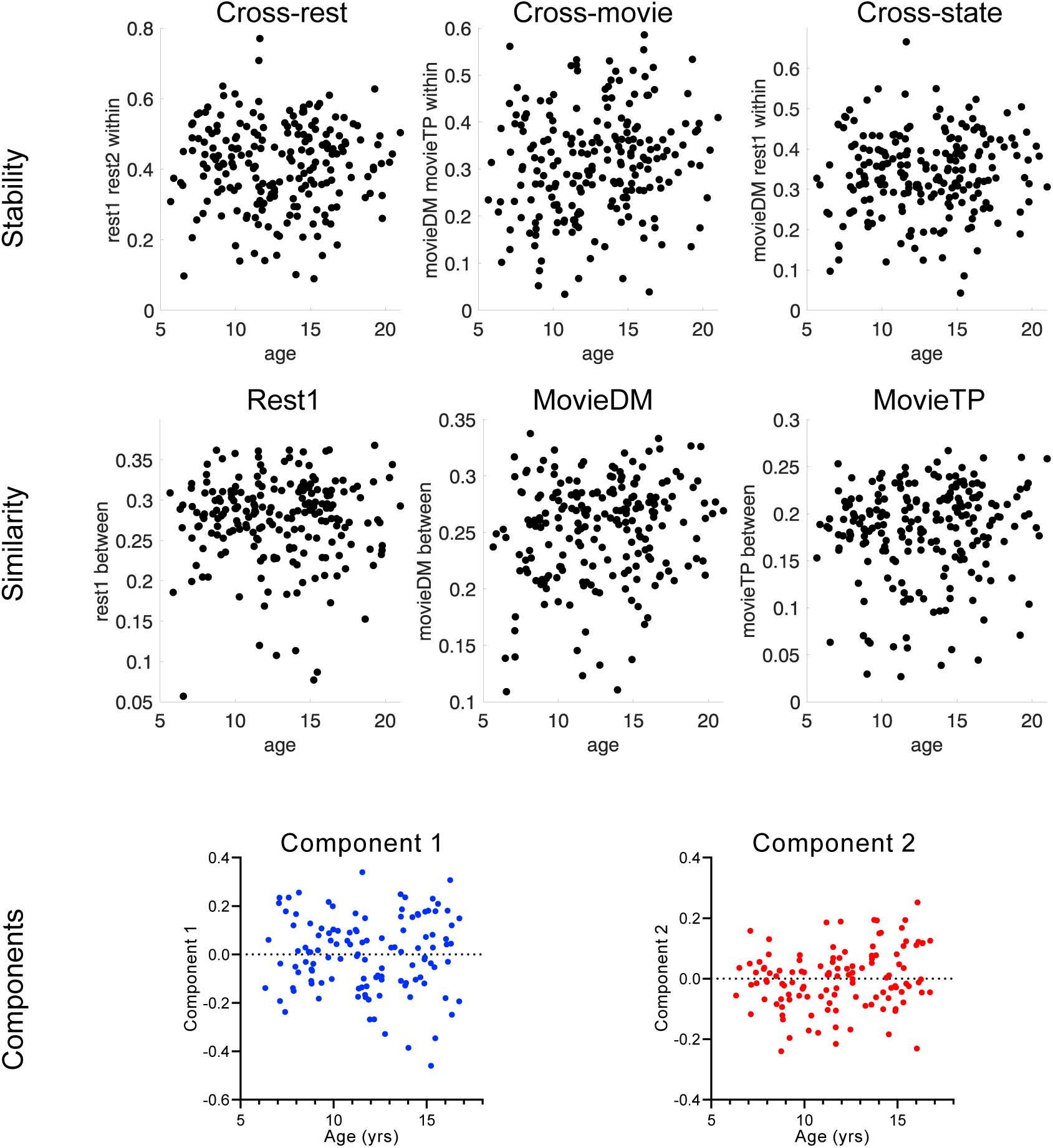

