## Supplemental Figures for "Stability and similarity of the pediatric connectome as developmental measures"

**Supplementary Materials**

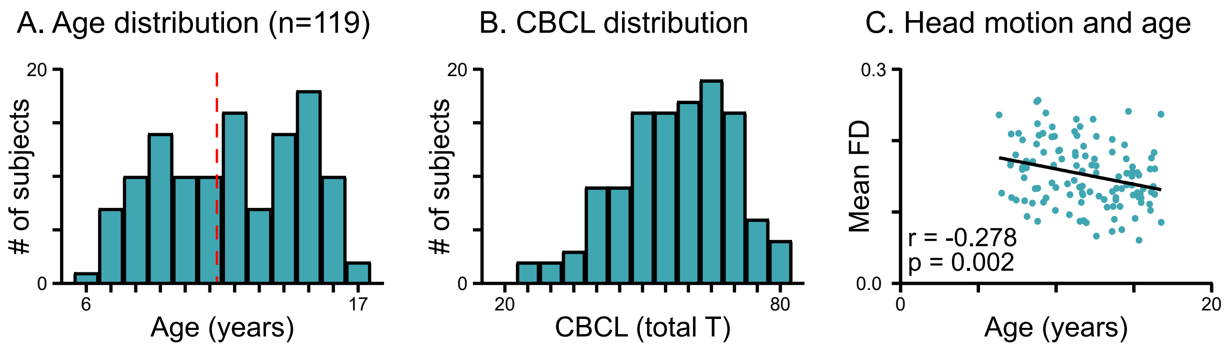


D. Behavioral measures of interest

| | CBCL total T | WISC-V FSIQ | SRS-2 total T |
|---|---|---|---|
| Age: r (p) | 0.112 (0.224) | -0.067 (0.467) | 0.071 (0.445) |
| Head motion: r (p) | 0.065 (0.485) | -0.001 (0.990) | 0.013 (0.891) |

**Supplementary Figure 1s: Demographics for behavioral subsample, n=119.** The age distribution, the right-skew of CBCL T-scores (in which greater scores indicate greater overall behavioral problems and psychopathology), and the association between advancing age and decreasing head motion are similar to the full 210 sample shown in Figure 1. D shows that there are no significant correlations among the three behavioral measures and either age or mean framewise displacement.

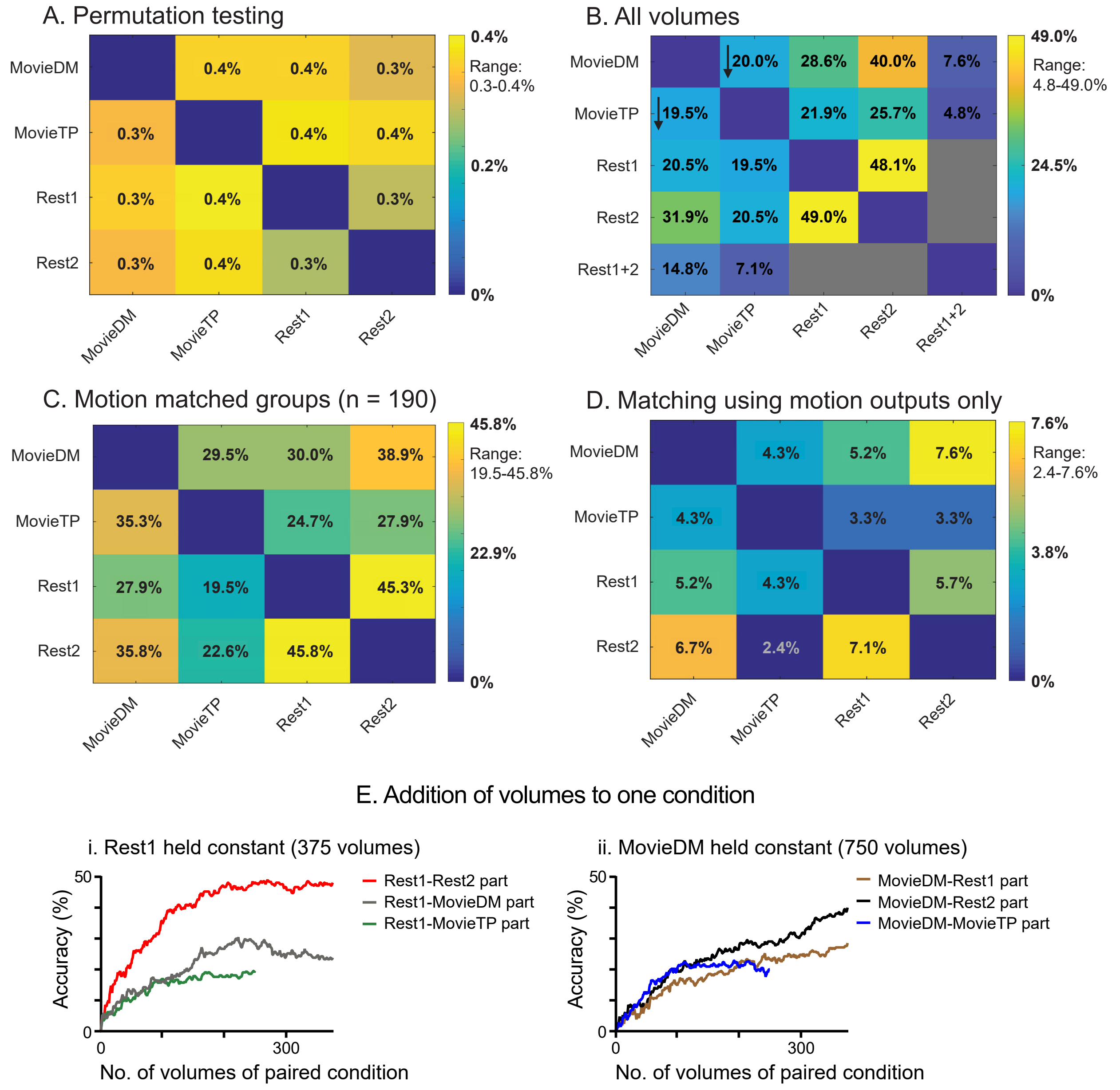


**Supplementary Figure 2s: Secondary test-retest matching algorithm analyses.** To further interrogate the results of the main matching algorithm reported in Figure 2, we conducted a number of other fingerprinting algorithms. **A)** Here we show the results of running the algorithm with permutations using falsely assigned identity pairs. **B)** The main analyses used matrices truncated at 250 volumes. Here, we reran the algorithm using all available volumes, so 750 volumes for MovieDM, and 375 volumes for Rest1 and Rest2. We also created an additional set of matrices, Rest1+2 (750 volumes), by concatenating across the two rest runs. The arrows indicate whether the values shown here are increased or decreased relative to the original analysis using thresholds set via permutation testing (p<0.05). **C)** There was a significant difference in mean framewise displacement across conditions, so here we reran the algorithm using a subset of subjects such that there was no difference in motion across conditions. All accuracies here are slightly higher than the original values. **D)** Here we reran the algorithm using only motion parameters. The graphs in **E)** show the effects of adding volumes to one matrix while holding the number of volumes in the other matrix constant. Contrary to expectations, adding data does not result in a linear increase in accuracy in all cases.

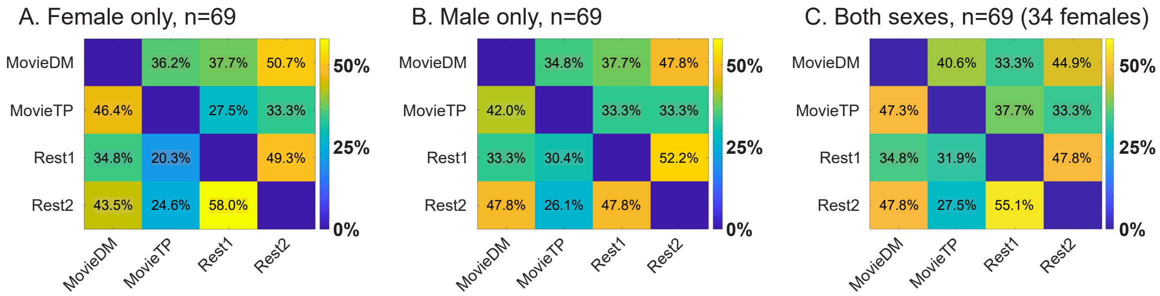


**Supplementary Figure 3s: Identification algorithms using single-sex cohorts.** Female- and male-only cohorts were identified that had no significant difference in mean age or mean FD, and the identification algorithms were rerun. Because sample size is a major determinant of accuracy, a mixed-sex cohort was also created to provide a comparison using the same sample size. All matrices show the same general pattern as observed in the full cohort (N=210) (Figure 2). We conducted permutation testing to assess whether cross-cohort differences were significant, and none met the permutation-based thresholds for significance ($p<0.05$). Neither cohort outperforms the others, and the overall increases in matching relative to the N=210 matrix are attributed to sample size. In this psychiatrically enriched, developmental cohort, using data from both sexes does not appear to have a major effect on matching accuracies of the test-retest identification algorithm.

A. Stability and similarity: correlations of all pairings

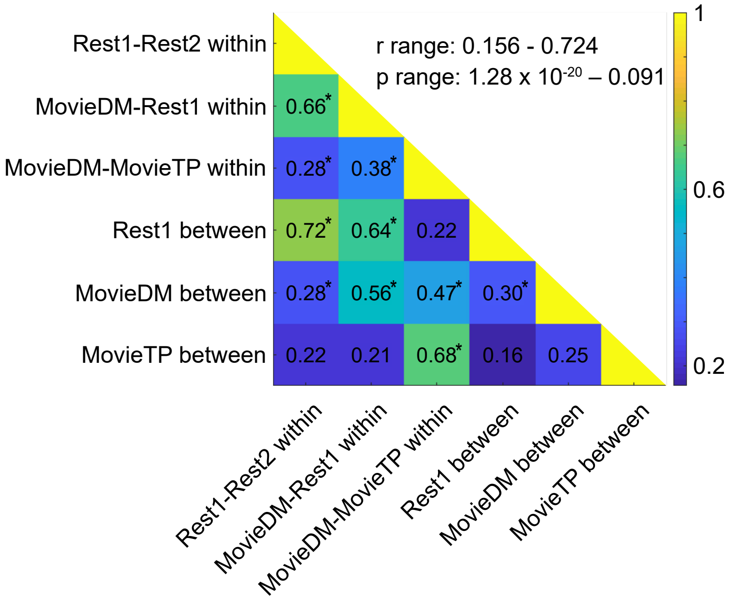


B. Stability and similarity: age and head motion

| | | Age: r (p) | FD: r (p) |
|---|---|---|---|
| Connectome Stability | cross-rest | 0.01 (0.91) | **-0.20 (4.13 x $10^{-3}$)** |
| | cross-movie | 0.15 (0.03) | **-0.38 (2.00 x $10^{-8}$)** |
| | cross-state | 0.07 (0.30) | **-0.29 (2.14 x $10^{-5}$)** |
| Connectome Similarity | Rest1 | 0.01 (0.89) | -0.15 (0.03) |
| | MovieDM | 0.16 (0.02) | **-0.41 (6.96 x $10^{-10}$)** |
| | MovieTP | 0.10 (0.15) | **-0.30 (8.53 x $10^{-6}$)** |

**Supplementary Figure 4s: All pairwise correlations among the six measures of connectome stability and similarity.** Main results for the stability/similarity relationships are shown in Figure 4, with head motion as a covariate. Here, we provide numerical values for all pairwise correlations of the six measures of stability and similarity (without covariates). Asterisks indicate statistical significance (Bonferroni corrected, $p<0.003$). Relationships of the six measures with framewise displacement (FD) and age are shown in B.

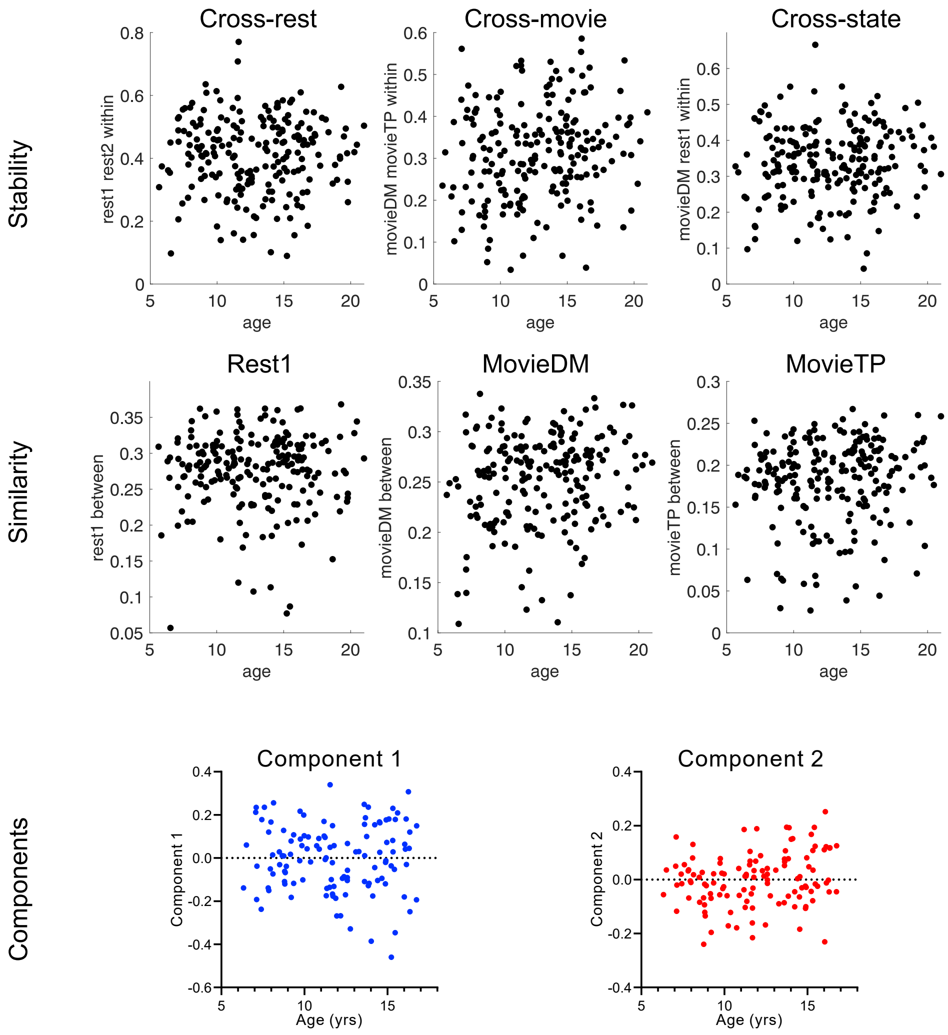


**Supplementary Figure 5s: Brain measures and age.** Neither the similarity and stability measures (N=210), nor the components from the PCA of the stability and similarity measures (n=119), show a clear relationship with age. When testing for a linear best fit, the only potentially significant correlation was between age and Component 2 ($r = 0.20$, $p=0.031$). When testing for a quadratic best fit, the only potentially significant correlation was also between age and Component 2 ($F(2,116)=3.54$, $r2 = 0.06$, $p=0.032$).
